# Excitatory TMS Boosts Memory Representations

**DOI:** 10.1101/279547

**Authors:** Wei-Chun Wang, Erik A. Wing, David L.K. Murphy, Bruce M. Luber, Sarah H. Lisanby, Roberto Cabeza, Simon W. Davis

## Abstract

Brain stimulation technologies have seen increasing application in basic science investigations, specifically towards the goal of improving memory functioning. However, proposals concerning the neural mechanisms underlying cognitive enhancement often rely on simplified notions of excitation and, most applications examining the effects of transcranial magnetic stimulation (TMS) on functional neuroimaging measures have been limited to univariate analyses of brain activity. We present here analyses using representational similarity analysis (RSA) and encoding-retrieval similarity (ERS) analysis in order to quantify the effect of TMS on memory representations. To test whether an increase in local excitability in PFC can have measurable influences on upstream representations in earlier temporal memory regions, we compared 1Hz and 5Hz stimulation to the left dorsolateral PFC. We found that 10 minutes of 5Hz rTMS, relative to 1Hz, had multiple effects on neural representations: 1) greater RSA during both encoding and retrieval, 2) greater ERS across all items, and, critically, 3) increasing ERS in MTL with increasing univariate activity in DLPFC, and greater functional connectivity for hits than misses between these regions. These results provide the first evidence of rTMS enhancing semantic representations and strengthen the idea that rTMS may affect the reinstatement of previously experienced events in upstream regions.

## Introduction

Over the past few decades, repetitive transcranial magnetic stimulation (rTMS) has developed into a powerful tool to establish causal brain-behavior relationships and improve cognition. Brain stimulation has been coupled with neuroimaging techniques in order to identify neural activity associated with enhancement of working memory (Mottaghy et al., 2002; Luber et al., 2007), episodic memory (Kohler et al., 2004), and semantic memory (Binney et al., 2010). However, most investigations on the effects of rTMS on neural substrates of memory functioning have been limited to univariate analyses of brain activity (Bollinger et al., 2010; Lee and D’Esposito, 2012; Vidal-Pineiro et al., 2014). Here we use a combination of rTMS, representational similarity analyses (RSA), and encoding-retrieval similarity (ERS) in order to test the efficacy of this technique in eliciting changes in neural representations that are a fundamental component of memory functioning.

Many rTMS studies have sought to relate various rTMS stimulation parameters to memory-related brain processes, focusing on the underlying neural correlates, rather than a measurable difference in memory. Because the relationship between any individual measure of memory and corresponding brain function is highly complex and is often mediated by multiple brain regions and psychological processes, this approach is advantageous for targeting specific components of neural function that contribute to memory encoding and retrieval. It is generally accepted that rTMS at higher frequencies (≥ 5Hz) induces local *increases* in BOLD activity during a host of different cognitive operations including motor planning (Schneider et al., 2010), working memory (Esslinger et al., 2014), and episodic memory (Vidal-Pineiro et al., 2014), *boosting* ongoing activity associated with successful performance. Conversely, 1Hz rTMS has been shown to reliably *depress* local hemodynamic activity (Muellbacher et al., 2000; de Vries et al., 2012; Plow et al., 2014). While the use of rTMS as a tool for behavioral neuroenhancement is undoubtedly informed by understanding these local effects, it remains unclear how local changes in neural activity affect large-scale system dynamics. Research combining rTMS with functional neuroimaging techniques has shown that rTMS can also have effects on distal brain regions that are functionally or structurally connected with the area targeted by stimulation (Esslinger et al., 2014; Wang and Voss, 2015). However, while the focus on “excitability” (a tenuous inference given the generality of the hemodynamic BOLD signal) in more distal regions of a circuit is becoming widespread (Fox et al., 2014), the effect of brain stimulation on the underlying representations upon which various cognitive processes operate has been almost completely unexplored.

To date, most applications examining the effects of rTMS on neuroimaging data have been limited to univariate analyses of brain activity. In contrast to standard univariate models, RSA instead asks how distributed patterns of brain activity evoked by different stimuli are related to one another, and thus provides the means to directly address questions of how mental representations are implemented in the brain as unique patterns of information. RSA methods have been used to explore how the relationship between stimuli of various dimensions—such as the visual or conceptual similarity of objects, words, or sentences—is reflected in the pattern of evoked neural responses associated with those stimuli. By comparing the similarity between each pair of stimuli in an fMRI dataset along a given dimension, we obtain a representational dissimilarity matrix (RDM), which serves to characterize a certain computational model of stimulus properties. These model RDMs can be compared to brain RDMs reflecting the similarity in activity patterns across the same pairs of stimuli. This approach has helped identify brain regions where representational structure captures higher-level semantic meaning that is not strongly contingent on stimulus modality (Devereux et al., 2013). Regions associated with semantic representations in RSA analyses includes left anterior temporal lobe (ATL) and left angular gyrus (AG) (Clarke and Tyler, 2014; Martin et al., 2018).

In the domain of episodic memory, representational approaches have been used to test a fundamental hypothesis that retrieving information from memory produces a partial recapitulation or reactivation of brain states and representations present when information was initially experienced (Danker and Anderson, 2010). One way of testing for this reactivation has involved comparing the similarity of brain patterns between the encoding and retrieval of items, known as encoding-retrieval similarity (ERS; Ritchey et al., 2013; Wing et al., 2015). Episodic memory reactivation has been reported in posterior cortical regions, as well as in the medial temporal lobes (MTL).

Whereas previous studies combining TMS with fMRI focused on univariate activity, we used multivariate RSA and ERS methods to investigate how TMS might alter the underlying stimulus representations at the time of initial learning, and the implications potential alteration might have for reactivated mnemonic information at retrieval. We applied TMS to DLPFC, a region which plays an important role in controlling relational memory binding (Blumenfeld and Ranganath, 2007), before a relational memory encoding-retrieval session in which fMRI data was collected. If DLPFC-focused stimulation is to have an effect in upstream locations, we expected that greater connectivity from the initial stimulation site to upstream sites managing these representations. Thus, we made three predictions: 5Hz relative to 1Hz TMS of DLPFC would enhance (1) the quality of semantic representations (as measured by RSA), (2) the quality of episodic memory representation reinstatement (as measured by ERS), and (3) DLPFC activity and functional connectivity accounting for the effect of rTMS on representations. Answers to these questions would help to clarify the variables that may influence the neural response to stimulation during a wide array of episodic memory paradigms.

## Materials & Methods

### Participants

Fourteen healthy older adults were recruited for this study (all native English speakers; 8 females; age mean +/- SD, 67.2 +/- 4.4 years; range 61-74 years); one subject was excluded to tolerability during rTMS, and two additional subjects were excluded due to not completing all imaging runs. Each older adult was screened for exclusion criteria for TMS (history of seizure, brain/head injuries) as well as psychiatric condition (MINI International Neuropsychiatric Interview, English Version 5.0.0 DSM-IV, (Sheehan et al., 1998)). None of the older participants reported subjective memory complaints in everyday life or had MMSE score below 27 (mean +/- SD = 29.1 +/- 0.8).

### Stimuli & Procedure

The associative memory task was comprised of a set of 360 sentences, each of which included a concrete subject and direct object. In each sentence, both subject and direct object were capitalized to indicate to the subject which specific nouns were to be remembered (*“A SURFBOARD was on top of the TRUCK.”*). Associative strength between nouns in a sentence (as determined by the USF word association norms (Nelson et al., 2004), were normally distributed, and both the imageability, frequency, and total length of each set of sentences was counterbalanced across all sentences used therein.

One run of the associative task comprised an encoding block of 90 sentences presented visually for 3 seconds, and a subsequent retrieval test comprising 68 word pairs from the same previously studied sentence, and 22 new pairs composed of two words recombined from two different sentences. Participants were asked to judge whether each word pair was intact or recombined and indicate how confident they were in their decision on a 4-point rating scale. Given the low proportion of low-confidence responses in the current data, we collapsed low and high-confidence responses, and excluded encoding trials in which subjects failed to indicate a response at retrieval.

### Image Acquisition

An outline of all data acquisition events is depicted in **Figure 1**. Scanning was divided between two days, 1-4 days apart, with the first day comprised of a functional memory-success localizer for scanning and stimulation on Day 2. All procedures were completed on a GE MR 750 3-Tesla scanner (General Electric 3.0 tesla Signa Excite HD short-bore scanner, equipped with an 8- channel head coil). Coplanar functional images were acquired with an 8-channel head coil using an inverse spiral sequence with the following imaging parameters: flip angle = 77°, TR = 2000ms, TE = 31ms, FOV = 24.0 mm^2^, and a slice thickness of 3.8mm, for 37 slices. The anatomical MRI was acquired using a 3D T1-weighted echo-planar sequence (256 × 256 matrix, TR = 12 ms, TE = 5 ms, FOV = 24 cm, 68 slices, 1.9 mm slice thickness). Scanner noise was reduced with earplugs and head motion was minimized with foam pads. Behavioral responses were recorded with a four-key fiber optic response box (Resonance Technology), and when necessary, vision was corrected using MRI-compatible lenses that matched the distance prescription used by the participant.

**Figure 1.**
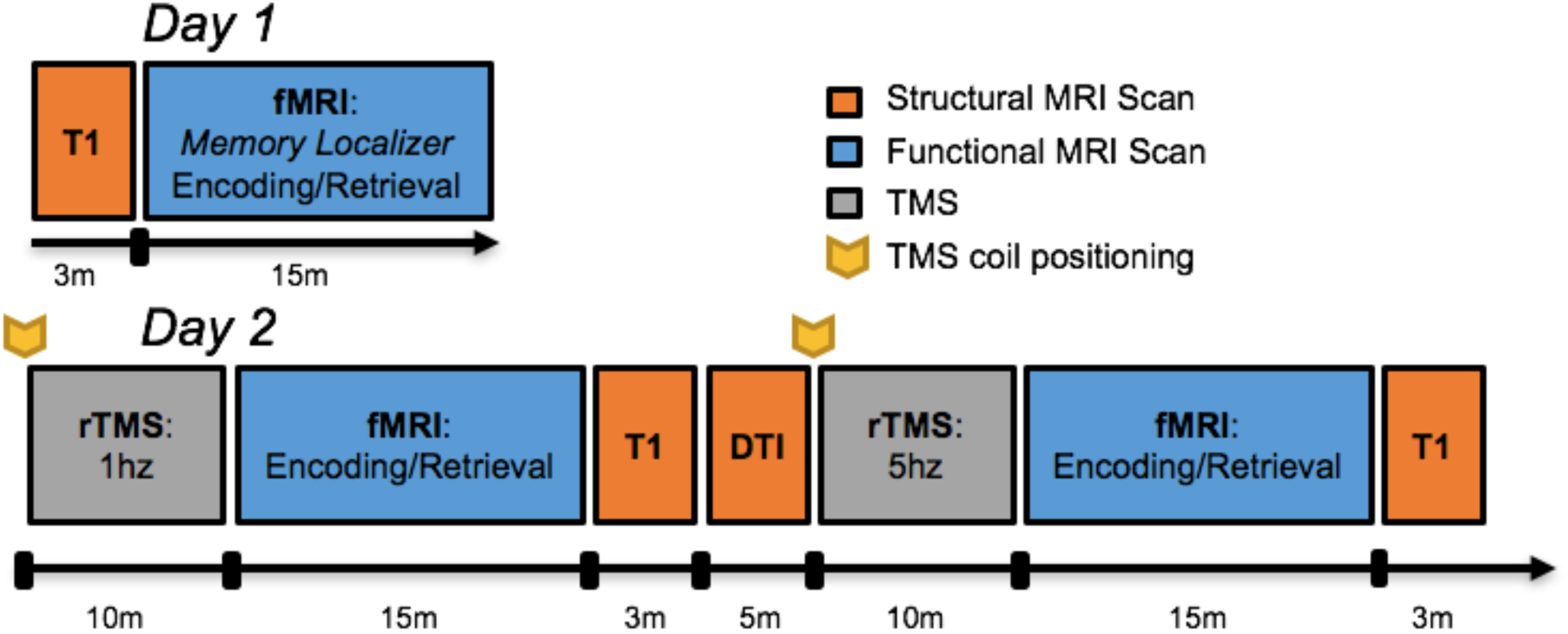
Timeline of imaging protocol.

### TMS Procedure

Prior to the TMS-fMRI session, the scalp location for TMS coil position over the left middle frontal gyrus (MFG) location had been found by using an infrared neuronavigation system (Brainsight: Rogue Research, Montreal, Canada). Specifically, the point of greatest activation in the left MFG in the fMRI memory contrast (i.e., encoding trials which were subsequently remembered versus forgotten) from the first day of scanning was chosen from the fMRI overlay on the subject’s structural MRI, both of which had been uploaded into BrainSight. After co-registration of the subject’s head with his MRI, the MFG location was marked on a tight fitting acrylic swim cap that stayed on the subject’s head until TMS-fMRI interleaving was completed on the same day. At that time, subjects were acclimated to the sort of TMS pulses to be delivered later in the scanner with a series of single pulses at the target site, as well as a short burst of 5Hz stimulation. The motor threshold (MT) for each subject was determined using a MagVenture R30M device located outside the scanner room, part of an MRI compatible TMS system which included a non-ferrous figure-8 coil with 12m long cable and artifact reducing counter-current charging system (MagVenture, Farum, Denmark). MTs were determined using electromyography of the right first dorsal interosseous (FDI) muscle and defined as the lowest setting of TMS device intensity at which ≥ 5 out of 10 motor evoked potentials of at least 50µV peak to peak amplitude could be elicited.

Before each functional scan, two 10-minute trains of either 1Hz or 5Hz stimulation were delivered at 120% MT immediately prior to an fMRI acquisition. The position of the TMS coil was reset to the same target site before the beginning of each rTMS session, and monitored continuously while the subject lay supine in the bed of the MR scanner. 1Hz rTMS was delivered in a continuous train of 10 minutes, while 5Hz rTMS was delivered in intermittent 6 sec trains with a 24 sec inter-train interval, also for 10 minutes. Dosage was equivalent between 1Hz and 5Hz rTMS conditions (600 total pulses), and the order of stimulation frequency was counterbalanced across subjects. Immediately after the 10 minutes of TMS, subjects were positioned in the scanner, and performed the encoding and subsequent retrieval portion of the sentence task while fMRI was acquired. The amount of time elapsed between the end of the rTMS train to the beginning of the functional scan was 9.4 minutes (SD = 1.7 minutes).

## Data Analyses

### Functional MRI preprocessing

Functional images were preprocessed using image processing tools, including FLIRT and FEAT also from FSL, in a publically available analysis pipeline developed by the Duke Brain Imaging and Analysis Center (https://wiki.biac.duke.edu/biac:analysis:resting_pipeline). Images were corrected for slice acquisition timing, motion, and linear trend; motion correction was performed using FSL’s MCFLIRT, and 6 motion parameters estimated from the step were they regressed out of each functional voxel using standard linear regression. Images were then temporally smoothed with a high-pass filter using a 190s cutoff, and normalized to the Montreal Neurological Institute (MNI) stereotaxic space. White matter and CSF signals were also removed from the data, using WM/CSF masks generated by FAST and regressed from the functional data using the same method as the motion parameters. Spatial filtering with a Gaussian kernel of full-width half-maximum (FWHM) of 6mm was applied.

To examine multivariate activity patterns, each fact was modeled in a separate GLM using the LSS approach on the unsmoothed data, yielding first-level single-trial beta images for each trial in each participant (Mumford et al., 2012) and were smoothed prior to group analyses. Group contrasts masked out white matter/CSF (SPM gray matter template values <0.1), all contrast images display effects at p <0.005, and cluster-level corrections for multiple comparisons to p <.05 with Monte Carlo Simulations (Slotnick et al., 2003; Slotnick, 2017).

### RDM Construction and RSA Analysis

Semantic similarity (RDMs) for the sentences (at encoding, **Figure 2A**) and the word pairs (at retrieval, **Figure 2B**) were constructed using the website cortical.io (Webber, 2015). Cortical.io captures the entirety of semantic space (trained on Wikipedia), distilled into a vector of 16,384 co-occurring words (i.e., ‘semantic contexts’). To calculate semantic similarity, it extracts the semantic contexts (in the vector of 16,384) associated with the words in each sentence at encoding and each word pair at retrieval (i.e., a binary ‘semantic fingerprint’ that is visualized with a 128^2^ matrix). To then create the RDMs, the dissimilarity (1-cosine similarity) between the semantic fingerprint for each pair of sentences or word pairs was then calculated. This method is relatively new, but has been successfully used to group similar firms based on their business descriptions (Ibriyamova et al., 2017) and group academic authors based on the content of their publications (Han et al., 2017). Thus, it provides a useful tool for extracting semantic similarity beyond just single words.

**Figure 2.**
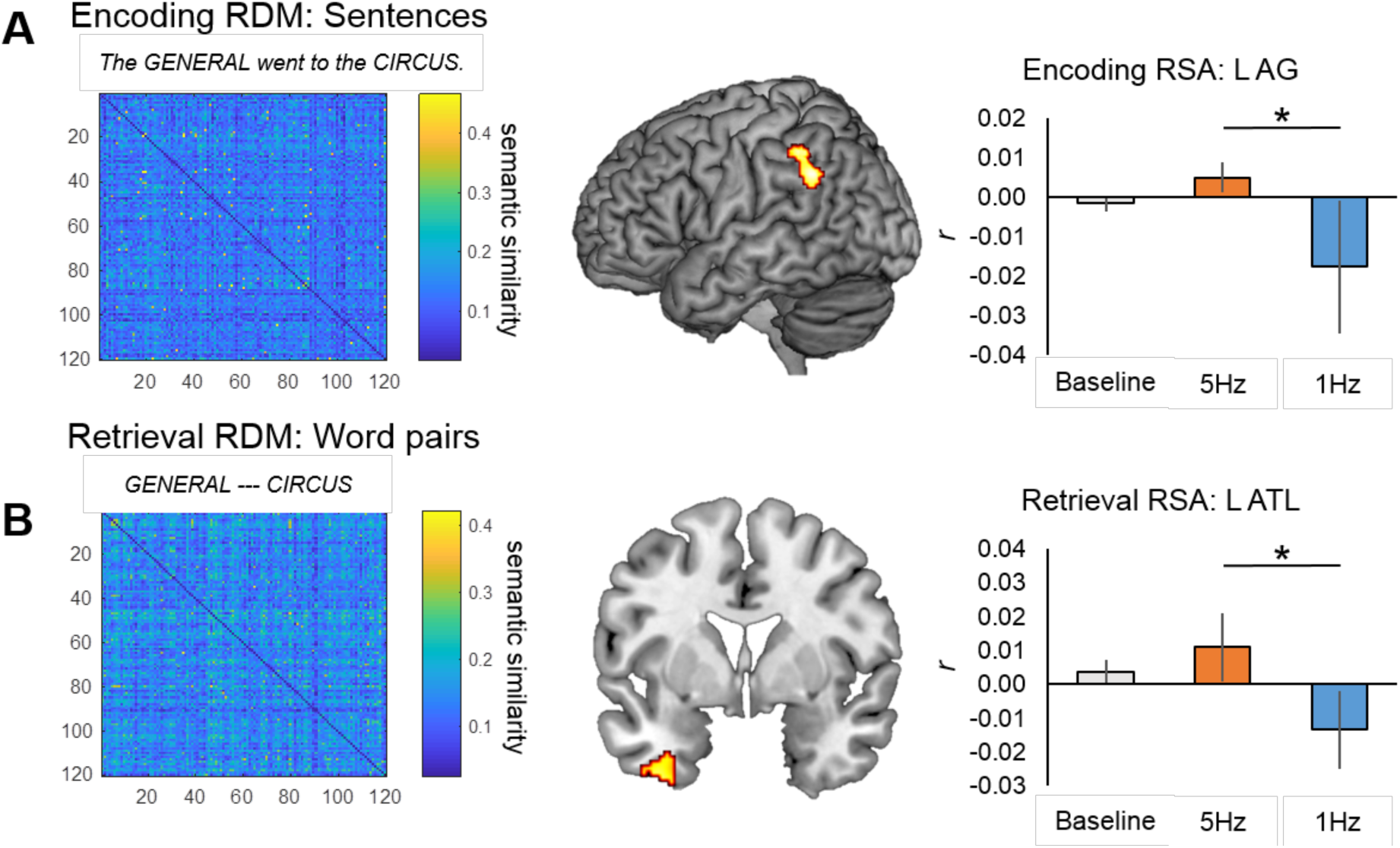
Regions exhibiting significant RSA effects during (**A**) encoding and (**B**) retrieval. Model RDMs based on sentential (**A**) or word pair similarity (**B**) are shown to the left, while at right are graphs representing mean 2^nd^ order correlations between model RDMs and brain RDMs.

**Figure 3.**
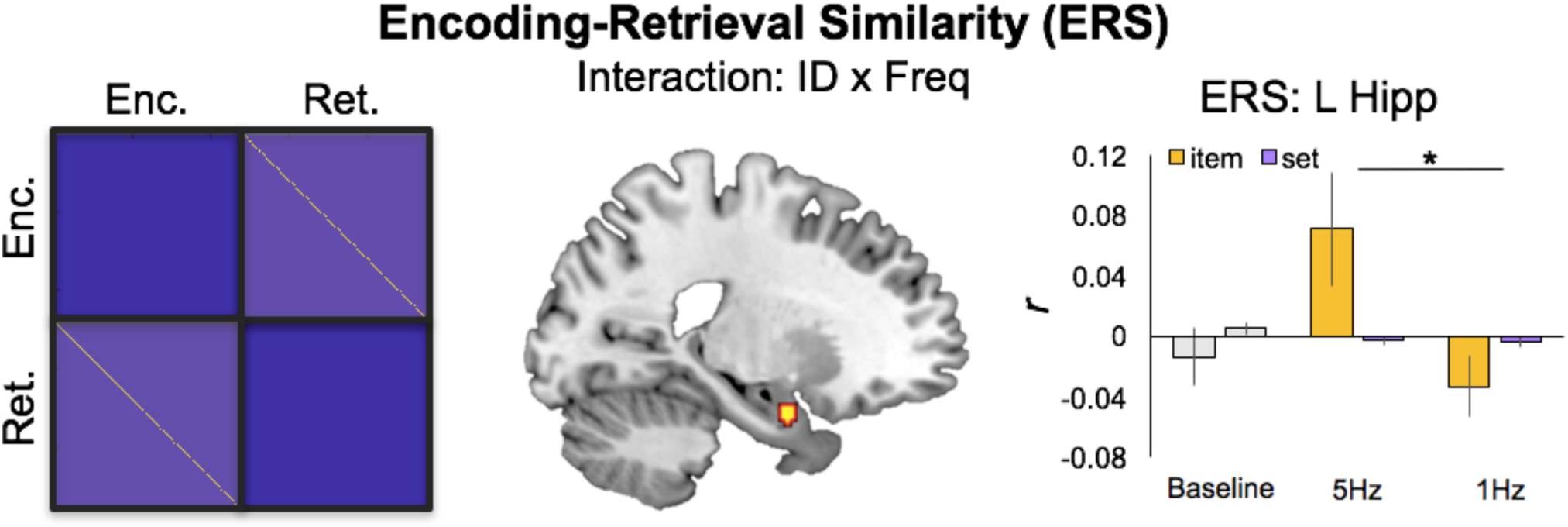
Encoding-Retrieval Similarity after 5Hz or 1Hz rTMS. The schematic on the left describes encoding-retrieval items to be compared (yellow cells) versus set-level matches between unmatched pairs (light purple cells). These set-level matches serve as a reasonable comparison for ERS effects. At right, mean pattern-to-pattern similarity across stimulation conditions for item- and set-level matches.

For each subject, second-order correlations were computed between our semantic RDM and single-trial beta images for each participant using an in-house searchlight script (https://github.com/brg015) with a 5-voxel searchlight sphere, separately for the three stimulation conditions. For group analyses, these second-order correlation maps were then spatially smoothed (6-mm isotropic FWHM Gaussian filter).

### ERS Analysis

ERS was calculated using an in-house searchlight script with a 5-voxel searchlight sphere (https://github.com/brg015) at both the item level, where encoding and retrieval trials involved the same item, and the set level, where encoding and retrieval trials belonged to the same set (i.e., condition) but involved different items. Item-ERS and set-ERS were calculated by correlating encoding and retrieval activity patterns within each searchlight sphere. For item-ERS, the encoding and retrieval activity patterns corresponding to the same trial were correlated, whereas for set-ERS, retrieval activity patterns for each trial was correlated with the encoding activity patterns for all trials in that stimulation condition (e.g., 5Hz) and then averaged to create the whole-brain similarity volume for that retrieval trial.

After ERS volumes had been calculated for each retrieval trail, fixed-effect contrasts were generated separately for item-ERS and set-ERS by averaging together all ERS volumes, yielding six mean ERS images per participant. For group analyses, these mean ERS images were then spatially smoothed (6-mm isotropic FWHM Gaussian filter). Because Baseline information was collected on a separate day, both RSA and ERS analyses are focused on the difference between RSA/ERS observed after two rTMS frequencies (5Hz, 1Hz); nonetheless, we include Baseline information here for descriptive purposes.

Lastly, to control for univariate activity, we also conducted confirmatory within-subject binary logistic regressions in clusters showing significant ERS effects. In this regression, stimulation condition (1Hz vs 5Hz) was the dependent variable, and there were three independent variables (IVs): (1) ERS, (2) univariate encoding activity, and (3) univariate retrieval activity. A test on the parameter estimates corresponding to the ERS regressor (see Table 3, rightmost column) indicated whether similarity measures uniquely predicted stimulation condition while accounting for the effects of univariate activity.

**Table 3.** Significant ERS clusters. Regression sig. (see methods): * p < .05 two tailed, † p < .05 one tailed

| ERS | *k* | *Z-score* | *x* | *y* | *z* | *Reg. t* |
| --- | --- | --- | --- | --- | --- | --- |
| Right Superior temporal gyrus | 114 | 3.49 | 48 | 0 | -18 | 2.71* |
| Left ATL | 43 | 3.08 | -40 | 0 | -26 | 3.00* |
| Left Hippocampus/Amygdala | 42 | 2.94 | -18 | 0 | -24 | 1.75† |

Note: Regression sig. (see methods): \* $p < .05$ two tailed, † $p < .05$ one tailed.

### Relationships between ERS and the DLPFC Stimulation Site

Lastly, we conducted two additional tests to examine the direct influence of left DLPFC stimulation on upstream representations. While the analyses above are focused on multivariate pattern information, estimates of univariate activity serve as useful markers of excitation or inhibition after 5Hz or 1Hz rTMS, respectively. Thus, we examined whether the magnitude of ERS correlated with this univariate activity in the region of stimulation in DLPFC. To do so, we extracted the mean univariate activity (beta values) in the DLPFC stimulation site, and ERS (Pearson’s *r* values) in clusters exhibiting significant effects and correlated them separately for each stimulation condition.

Second, we used task-based functional connectivity estimates to determine the degree to which these regions were coupled with DLPFC across encoding, i.e., immediately after 5Hz or 1Hz rTMS. Functional connection matrices representing task-related connection strengths were estimated using a correlational psychophysical interaction (cPPI) analysis (Fornito et al., 2012). Briefly, the model relies on the calculation of a PPI regressor for each region, based on the product of that region’s timecourse and a task regressor of interest, in order to generate a term reflecting the psychophysical interaction between the seed region’s activity and the specified experimental manipulation. In the current study the task regressors based on the convolved task regressors from the univariate model described above were used as the psychological regressor, which coded subsequently remembered and subsequently forgotten word pairs with positive and negative weights, respectively, of equal value. This psychological regressor was multiplied with two network timecourses for region *i* and *j*. We then computed the partial correlation *ρ_ppI_i__*,*_ppI_j._z_*, removing the variance *Z* associated with the psychological regressor, the timecourses for regions *i* and *j*, and constituent noise regressors. We accounted for the potential effects of head motion and other confounds by assessing the 6 motion parameters and including these parameters in our partial correlation between regions.

## Results

### Behavioral results

As explained in the Introduction, the goal of the current analysis was to investigate the effects of TMS on mnemonic representations and their fidelity across encoding and retrieval conditions. Behavioral results for our associative memory task are summarized in **Table 1**. We found no significant differences between rTMS conditions, either in Hit Rate (t(10) = 1.55, p = 0.15), False Alarm Rate (t(10) = 1.01, p = 0.34), or d’ (t(10) = 0.37, p = 0.72); response times for correct trials were similarly (all p > 0.05). These null findings for behavioral differences between rTMS conditions suggest that any brain-related differences in multivoxel pattern information (RSA/ERS) are not readily attributable to differences in task strategy or strength of encoding.

**Table 1.** Behavioral performance

| *Accuracy* | H (SE) |  | FA (SE) |  | d' (SE) |  |
| --- | --- | --- | --- | --- | --- | --- |
| Baseline | 0.77 | 0.04 | 0.15 | 0.04 | 2.01 | 0.24 |
| 1Hz rTMS | 0.82 | 0.05 | 0.11 | 0.03 | 2.35 | 0.21 |
| 5Hz rTMS | 0.87 | 0.03 | 0.14 | 0.03 | 2.42 | 0.23 |

| *Response Time* | H (SE) |  | FA (SE) |  |
| --- | --- | --- | --- | --- |
| Baseline | 1814 | 107 | 2450 | 111 |
| 1Hz rTMS | 1687 | 99 | 2410 | 112 |
| 5Hz rTMS | 1523 | 102 | 2143 | 131 |

### RSA results

Our main goal was to examine evidence that rTMS has upstream effects on the quality of information processed in a standard memory paradigm. To test our first prediction on the effects of TMS on semantic representations, we examined the influence of rTMS on the semantic quality of representations (2^nd^ order correlation with the semantic RDM) during the Encoding period. The searchlight RSA revealed greater second-order correlations between activity patterns and our semantic RDM for excitatory (5Hz) than inhibitory (1Hz) rTMS in posterior regions including the left AG, bordering the supramarginal gyrus (**Figure 2A**; **Table 2**, top panel). No significant clusters emerged in the reverse contrast (i.e., 1Hz > 5Hz). Next, we examined the fidelity of representations during the Retrieval period, when participants were completing an associative recognition test for word pairs from the sentences read at encoding. The searchlight RSA revealed greater second-order correlations between activity patterns and our semantic RDM for 5Hz than 1Hz rTMS in regions including the PHC and ATL (**Figure 2B**; **Table 2**, bottom panel). Again, no clusters emerged in the reverse contrast (1Hz > 5Hz). In sum, consistent with our first prediction, we found that TMS enhanced the quality of semantic representations (greater 2^nd^ order correlation with semantic RDM) in typical semantic regions such as left AG and left ATL.

**Table 2.** Significant RSA clusters.

| Encoding RSA |  |  |  |  |  |
| --- | --- | --- | --- | --- | --- |
| Region | k | Z-score | x | y | z |
| Left AG/SMG | 68 | 3.22 | -62 | -50 | 34 |
| Right AG | 31 | 3.03 | 56 | -54 | 20 |
| Cuneus | 36 | 2.95 | -16 | -74 | 22 |
| Retrieval RSA |  |  |  |  |  |
|  | k | Z-score | x | y | z |
| Left Precentral gyrus | 111 | 3.73 | -16 | -24 | 66 |
| Left Anterior PHC | 92 | 3.59 | 32 | -22 | -16 |
| Left ATL | 148 | 3.30 | -38 | 4 | -44 |
| Posterior cingulate | 24 | 3.01 | 12 | -16 | 38 |

### ERS Results

To test our second prediction, we then examined the influence of rTMS on the fidelity of memory representations measured using ERS. ERS allows us to identify the reactivation of episodic representations between encoding and retrieval by comparing activity patterns at encoding and retrieval for trials belonging to the same item (i.e., item-ERS) and trials belonging to different items (i.e., set-ERS). We examined whether any regions exhibited stimulation by ERS level interactions. This contrast revealed greater item-ERS than set-ERS for 5Hz than 1Hz stimulation in MTL and temporal regions (**Table 3**). These effects were significant even when controlling for univariate activity (**Table 3**, rightmost column). No clusters exhibited the opposite interaction (i.e., 1Hz > 5Hz). Thus, greater item-than set-level ERS helps to confirm our second prediction, such that excitatory 5Hz rTMS to DLPFC enhanced episodic memory representations in MTL.

### Relationships between ERS and the DLPFCLPFC Stimulation Site

Lastly, to test our third prediction that the representational would be associated with increases in DLPFC activity and connectivity. We first extracted univariate activity at the stimulation site during encoding (**Figure 4A**). As reported in our previous analysis of these data (Davis et al., 2017), the univariate activity for successfully remembered trials in the stimulated left MFG region was reduced by 1 Hz rTMS compared with Baseline (t = -2.43, P < 0.01), but was increased by 5 Hz rTMS (t = 2.83, P < 0.01) compared with Baseline activation for subsequently remembered encoding trials. A pairwise comparison between 1Hz and 5Hz fMRI activity for successfully remembered trials was also significant (t = 3.43, P < 0.001). This result confirms that rTMS modulated memory-related activity at our site of stimulation and enables our two subsequent analysis of how this upregulation of univariate activity in left PFC is related to upstream effects in MTL.

**Figure 4.**
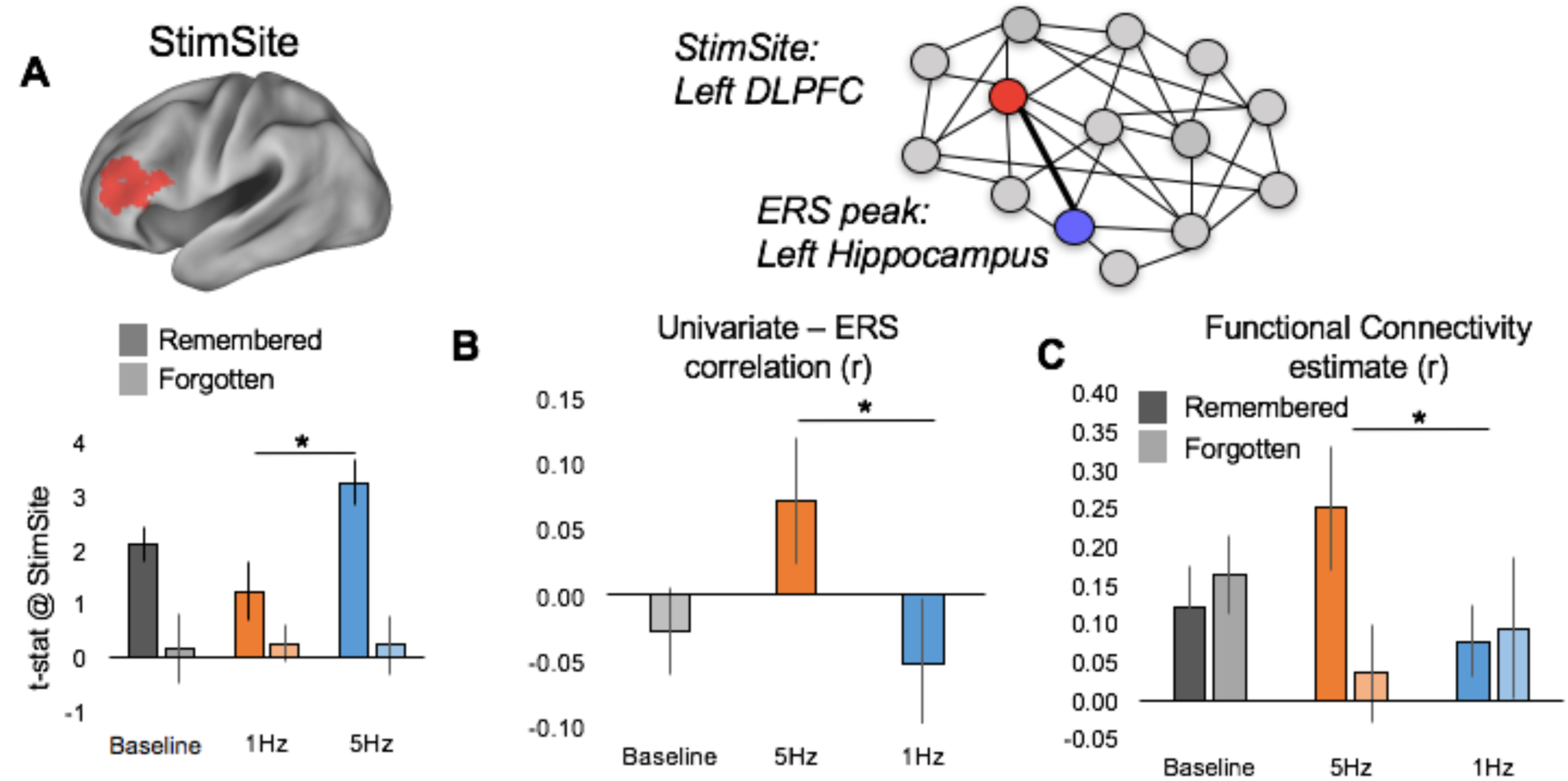
Relationships between Left PFC stimulation site and MTL ERS effects. (**A**) The left PFC Stimulation Site within the left MFG is shown in red, with standard effects for subsequently Remembered and Forgotten trials during Baseline, 1Hz, and 5Hz conditions. (**B**) This increase in univariate activity served to boost ERS in left hippocampus after 5Hz rTMS. (**C**) Functional connectivity estimates based on the cPPI analysis during the encoding period, which demonstrate a selective increase in PFC-MTL connectivity during successfully remembered > forgotten trials, and effect present only in the 5Hz rTMS condition.

We then examined whether univariate activity at the stimulation site (DLPFC) correlated, across trials, with the magnitude of our observed stimulation by ERS level interaction effects. This analysis found a significantly greater correlation (t(10) = 2.28, p < .05) between left DLPFC activity at encoding and the left MTL cluster during 5Hz stimulation than 1Hz stimulation (**Figure 4B**). There were no significant correlations found in any other ERS regions.

Lastly, in order to link local stimulation effects in the DLPFC to representational changes as a function of memory success, we examined the functional connectivity between the DLPFC stimulation site and regions exhibiting greater ERS following excitatory stimulation. We found a significant Success x Frequency interaction (F = 3.76, p = 0.04), wherein connectivity at encoding between DLPFC and the left MTL cluster was greater for subsequently remembered than forgotten trials following 5Hz compared to 1Hz rTMS (**Figure 4C**). There were no significant correlations found in any other ERS regions. These results help to clarify the role of DLPFC in mediating upstream semantic representations that are important for successful memory functioning.

## Discussion

To our knowledge, this is the first study to demonstrate that rTMS can boost memory representations as measured by RSA and ERS. Thus, this work links brain stimulation techniques with semantic representations and episodic memory functioning using a novel multimodal paradigm. The study yielded three principal findings. First, 5Hz, relative to 1Hz, rTMS boosted the quality of semantic representation (i.e., 2^nd^ order correlation with semantic RDM) in left AG and left ATL. Second, 5Hz, but not 1Hz, rTMS enhanced quality of episodic memory representations in MTL, as assessed by ERS analyses. Lastly, linking the ERS effect to the DLPFC stimulation, we found that increases in univariate activity at the stimulation site were correlated with increases in item-level ERS in MTL, and that these two sites were more functionally connected during subsequently remembered trials. We discuss each of these results and their implications below.

### rTMS effects on Semantic Representations

Our result demonstrates that external manipulations of local excitability in left DLPFC impact neural pattern similarity in distinct temporal and parietal regions known to code for semantic and episodic features. Foundational to this finding is the idea that the neural and semantic similarity between individual items represents meaningful information, and that this representation in a given brain region or computational model can be characterized by the matrix of dissimilarities between the stimulus representations (Kriegeskorte and Kievit, 2013). The information encoded in our task represented complex sentences with conceptual memoranda common to everyday life. Despite its fundamental role in cognition, the informational content that is represented during reading remains elusive. To understand how semantic information is represented in the brain, we used a comprehensive database of relationships between semantic contexts capturing a broad semantic space (Cortical.io). The similarity between the semantic contexts in each sentence (encoding) or word pair (retrieval) across our task can be summarized with a single representational dissimilarity matrix (or RDM), and thus presents a powerful means of describing relatively complex information patterns across multiple trials.

While no study has used TMS to examine changes in pattern similarity, a number of studies have targeted activation patterns associated with more upstream components of conceptual encoding. Recent studies suggest that TMS, when applied early in the ventral processing stream, may strongly impact univariate estimates related to visual and mnemonic processing. 20Hz TMS applied to early visual cortex during short-term retention of visual stimuli results in a reduction in behavioral measures of working memory precision (van de Ven et al., 2012; van de Ven and Sack, 2013), supporting the notion that visual cortex stores precise representations of visual working memory contents. While most of these studies are focused on disrupting posterior regions that are thought to represent stimulus-specific information, there is also evidence that disruption of semantic processing can also occur as a consequence of 1Hz rTMS to the anterior temporal lobes (Lambon Ralph et al., 2009; Binney et al., 2010).

Our results provide a clear step forward for the investigation of the encoding and retrieval of semantic concepts. Critically, both RSA and ERS were greater for trials after excitatory stimulation in regions typically associated with more abstract semantic representations like the ATL and AG. These results suggest that conceptual information represented in in temporal and parietal regions is modulated by TMS. For meaningful word stimuli, recent work has shown that representations in ATL relate to measures of true and false memory, suggesting that exogenously induced representational changes in this region might underpin aspects of memory function for similar stimuli (Chadwick et al., 2016). These convergent results therefore suggest a distinct mechanism whereby successful memories become consolidated and available for remembering through specific neural patterns. To date there is no evidence that brain stimulation can modulate these effects, but the current results help to shed some light on this potential mnemonic process.

### Top-down influences on memory

As noted in the Introduction, one needs to consider that the region-specific effect of rTMS on memory performance might have been boosted by indirect manipulation of regions connected with the stimulation site. The current results provide preliminary evidence that excitatory influences in DLPFC can have meaningful influences on the representational information processed in more posterior cortices. We found two complimentary pieces of evidence to support the idea that PFC-mediated increases in representational similarity can be induced with rTMS. First, and most critically, the local activity increases at the site of stimulation were more strongly correlated with item-level ERS in the left MTL during excitatory relative to inhibitory stimulation. Second, functional connectivity between the DLPFC stimulation site and the left MTL during encoding demonstrated a significant Stimulation x Memory Success interaction (**Figure 4C**), such that connectivity was greater during successfully encoded trials than subsequently forgotten trials after 5Hz (but not 1Hz) stimulation. These results suggest that representations in MTL were boosted by indirect manipulation.

While this is, to our knowledge, the first time TMS has been shown to modulate pattern similarity, it is not the first to suggest that PFC stimulation may boost the top-down activation (or controlled retrieval) of semantic knowledge (Badre and Wagner, 2007; Bunge et al., 2009). Causal links between PFC activity and the properties of visual cortical neurons has been established through intervention with TMS. Disruption of PFC activity using 1Hz rTMS causes a significant reduction in the selectivity of fMRI responses in visual cortex, and appears to inhibit top-down selective attentional modulation to specific category types in the occipitotemporal cortex, during both online perception (Higo et al., 2011), and during working memory delay phases, (Lee and D’Esposito, 2012).The principle implied here is that DLPFC inputs enhance selectivity in visual cortex; given that our stimulation region was more functionally connected to MTL regions exhibiting significant ERS effects, our results extend these findings to regions implicated in semantic memory and go beyond univariate measures in suggesting that the representational content evoked during encoding and reactivated during retrieval is fundamentally affected by stimulation in frontal control regions.

### The importance of frequency

Lastly, it is key to note that both our RSA/ERS pattern similarity findings, and that success-related functional connectivity between left DLPFC and left MTL during encoding (**Figure 4C**) were uniquely associated with fMRI data collected after 5Hz rTMS. This result is reasonable given the importance of frontotemporal synchrony within the theta band (4-10Hz) during episodic encoding in regions functionally connected to the hippocampus, including LPFC and VPC (Osipova et al., 2006; Hsieh and Ranganath, 2014). EEG signals generated by the hippocampus are typically dominated by regular theta waves, often continuing for many seconds after being elicited by a focused event (Buzsaki, 2002, 2005).

Increased rhythmic synchrony across distant regions within a cortical network is thought to improve information processing by increasing network efficiency, an effect particularly important during demanding cognitive processing (Fries, 2009; Deco et al., 2011), such as episodic encoding and retrieval. For example, a number of scalp EEG studies have reported that frontotemporal theta rhythms are enhanced for items that are subsequently remembered more than items that are forgotten (Summerfield and Mangels, 2005; Hanslmayr et al., 2009; Nyhus and Curran, 2010). Even more recent intracranial work from Kahana and colleagues has helped to validate this observation, which suggest that widespread synchronization of theta networks predict successful encoding (Ezzyat et al., 2017; Solomon et al., 2017), and that selective modulations of theta- and high-gamma-band activity in the temporal lobes can be used to decode successful memory states and inform targeted, closed-loop brain stimulation paradigms (Ezzyat et al., 2018). Such direct applications of frequency-specific information and brain stimulation represent powerful tools for future research and clinical application.

### Caveats

Our study has a number of limitations. First, there are unique subjective differences between the experience of 1Hz and 5Hz rTMS. Thus, it is possible that greater RSA/ERS effects after 5Hz TMS were due to subjects being cued by the greater intensity of sound produced by a 5Hz rTMS and may have been in a state of heightened awareness. While we cannot discount this possibility, we note that 1) the memory task was not concurrent with the rTMS, so the influence of such heightened attention would have to persist through the ∼10 minutes necessary to initiate the fMRI scan, 2) we observed no significant univariate effects of 1Hz stimulation compared to baseline, suggesting that the application of any TMS was not sufficient to elicit this effect, and 3) performance on the memory task did not differ as a function of stimulation frequency.

Second, our study suffers from a low sample size; a recent meta-analysis of clinical rTMS studies found that the average sample size for such studies is low (mean sample size = 18; Martin et al., 2003). Clearly this standard is not viable for the field to progress beyond qualitative observations of rTMS impact. However, our findings support the conclusion that the modulatory influence of rTMS may be limited to specific frequency parameters, offer an exploratory foundation in characterizing representational changes associated with rTMS, and suggest an analytical format that may reveal important information as yet unexplored in more well-powered brain stimulation studies.

## Conclusion

Overall, our findings indicate a direct link between 5Hz rTMS applied to DLPFC, and memory representations in posterior parietal and temporal cortices. Taken together, these results provide the first evidence of excitatory TMS enhancing memory representations using two different multivariate methods and suggests that rTMS may affect the reinstatement of previously experienced events in regions distal to the site of activation. By showing that stimulation impacts the neural substrates of episodic memory at specific frequencies, our data provide a foundation for future work to apply stimulation when it is most likely to improve memory function.

